# Feeding the host reshapes virulence: nonlinear scaling in a microsporidian pathogen

**DOI:** 10.64898/2026.03.26.714583

**Authors:** Charlotte Carrier-Belleau, Michael Officer, Niamh McCartan, Joseph Strawbridge, Nurfatin Zulkipli, Jeremy J. Piggott, Pepijn Luijckx

**Author notes:** Equal contribution (Charlotte Carrier-Belleau, Michael Officer).

## Abstract

Resource availability is a central driver of ecological and evolutionary processes, yet its effects on infectious disease and virulence are not fully understood. A key limitation is that many studies consider only a narrow range of resource conditions or a limited subset of host and pathogen traits, potentially obscuring non-linear relationships. Here, we quantify how a gradient of six food levels simultaneously shapes host fitness and pathogen performance in the *Daphnia magna*– *Ordospora colligata* system. Across two laboratory experiments, we measured infection rates, pathogen burden, host fecundity, survival, and filtration rates. Increased food availability enhanced pathogen fitness, with both infection rates and spore burden increasing with provisioning. In contrast, host responses were trait-specific: while fecundity increased with food availability, pathogen-induced reductions in fecundity (i.e., virulence) peaked at intermediate resource levels, despite continued increases in pathogen load. This pattern indicates that resource availability alters host tolerance as well as pathogen growth, generating non-linear disease outcomes. Host survival was unaffected by either food provisioning or infection, further demonstrating that resource availability can simultaneously influence host and pathogen traits in different directions. Our results highlight the importance of integrating multiple fitness components across provisioning levels to understand disease dynamics and suggest that ongoing anthropogenic changes in resource availability may alter host–pathogen interactions.

## Introduction

Resource availability is a fundamental ecological driver that influences individual fitness (Schälicke et al., 2019), population growth (Prevedello et al., 2013), and community dynamics (Galbraith et al., 2015). Variation in food availability can shape life history traits like age of maturity (Agrelius et al., 2023), affect immunity (Marcos et al., 2003), alter behaviour (e.g., mating tactics (Kolluru and Grether, 2005), or migratory patterns (Whelan et al., 2020)), and affect the outcome of species interactions. Indeed, impacts of resource availability on both predator–prey and host– pathogen systems have been reported (Duffy et al., 2012; Rosenzweig, 1971; Tadiri et al., 2013). For example, when resources were supplemented in a stream, large-bodied consumers increased, but predators did not, as they were unable to capture the larger prey (Davis et al., 2010). Similarly, in host–pathogen systems, increased food availability can enhance host growth and reproduction, but may also alter infection: well-nourished hosts can mount stronger immune responses and resist pathogens, whereas in some cases, increased provisioning can facilitate parasite transmission by promoting host aggregation or higher host density (Becker et al., 2015). Indeed, resource availability can impact both animal and human health by modifying disease dynamics (Tadiri et al., 2013; Weger-Lucarelli et al., 2018). With human-driven resource supplementation through agricultural activities, waste disposal, and the provision of supplementary food sources (e.g., bird feeders, crop residues) and global change altering productivity (O’Connor et al., 2009), it is thus especially important to understand how this alters species interactions. Moreover, food supplementation in disease systems may lead to impacts distinct from those observed under predation or competition (Cressler et al., 2014).

The relation between resource availability and disease outbreak and virulence is complex, and can be influenced by numerous factors (Cressler et al., 2014; Pike et al., 2019). An increase in resources can elevate population density or may lead to aggregation of hosts in areas where resources are provided (Civitello et al., 2018), leading to greater contact rates and thus enhanced disease spread. Improved nutrition to the host can also enhance pathogen growth as more resources are available for replication (Bedhomme et al., 2004). However, whether increased food provision leads to increased pathogen load and virulence may depend on how the pathogen and the host immune system compete for those resources (Cressler et al., 2014). While an increase in pathogen load is predicted when the pathogen is not competing with the immune system for resources, pathogen load may decrease if the immune system can monopolize or restrict the resources. Indeed, hosts may reduce the availability of glucose in response to infection (Kreimendahl and Pernas, 2024), as seen in mice infected with *Plasmodium* (Ramos et al., 2022). Alternatively, if both the pathogen and the immune system compete for the same resources, pathogen load may peak at intermediate levels of food supplementation (Cressler et al., 2014). Outcomes may be further complicated when resource availability alters host behaviour (Becker et al., 2015) or if virulence is in part caused by self-damage done by the immune system, which could be higher when animals are well-fed (Cornet et al., 2014). For example, altered feeding behaviour of the crustacean *Daphnia* under high food conditions leads to reduced filtration rates (Chow-Fraser and Sprules, 1992) and thus lower encounter rates with pathogens (Hall et al., 2007). Indeed, numerous studies show that the interplay between pathogens, hosts, and food provisioning can lead to context- and system-specific outcomes (Becker et al., 2015; Cressler et al., 2014) in part because multiple host and pathogen traits might be differentially impacted (Borer et al., 2023). There is thus an urgent need for more studies to disentangle how food provision shapes within-host growth of pathogens, virulence (Pike et al., 2019) and disease dynamics (Borer et al., 2023)

Here, we studied the effect of increased availability of resources on the crustacean *Daphnia magna* infected with the microsporidian pathogen *Ordospora colligata*. This host-pathogen system has been used to understand how rising temperatures and extreme weather alter disease proliferation (Kirk et al., 2018; McCartan et al., 2025), and the outcome of multiple infections (O’Keeffe et al., 2024). These studies have reported a wide range of pathogen loads, and while this pathogen has initially been described as benign (Ebert, 2005), reductions in host fecundity of up to 60% have recently been reported (O’Keeffe et al., 2024). Given that these studies used variable amounts and species of algae as food for the host, this may explain the variability in the observed responses. Meta-analysis suggests that in invertebrates (Cressler et al., 2014), like *Daphnia*, increased food provisioning often leads to increased pathogen loads, which could have a greater impact on their host, although not necessarily so, as increased nutrition may also increase tolerance (Budischak and Cressler, 2018; Miller and Cotter, 2018). Indeed, the outcome may be complex as increased food provisioning could lead to decreased contact if *Daphnia* filter less water when food is abundant (Chow-Fraser and Sprules, 1992) or if tolerance to disease is highest at lower levels of food provisioning as reported for *Drosophila* infected with *Salmonella typhimurium* (Ayres and Schneider, 2009). Here, we conducted two laboratory experiments with *Daphnia* and its gut pathogen *O. colligata* across a food gradient. Because many studies use only a limited number of food levels or measure only one or a few host and pathogen traits, non-linear patterns and other complexities in host and pathogen performance may remain undetected. (Borer et al., 2023). Our design that incorporates six levels of food provisioning allows us to assess how multiple host and pathogen fitness components scale with food provisioning, revealing non-linearities and their implications for virulence.

### Study System

The freshwater crustacean *D. magna* has been widely studied as a model organism for ecological and evolutionary processes (Ebert, 2022, 2005). As a filter feeder of planktonic algae and other microorganisms, *Daphnia* play a key role in freshwater ecosystems (Ebert, 2005). Their short generation time, ability to reproduce both sexually and asexually - which allows diverse clonal lines to be maintained - and availability of numerous pathogens have also made them widely studied as a system explaining host-pathogen coevolution (Decaestecker et al., 2007), disease dynamics (Duffy et al., 2012), and the outcome of multiple infection (O’Keeffe et al., 2024). Here we use *O. colligata* a microsporidium parasite that exclusively infects *D. magna.* Spores of this gut pathogen are released into the environment when infected gut cells lyse and are expelled with the faeces. Hosts encounter these spores through filter feeding. Once ingested, the spores penetrate cell membranes in the upper gut, where they develop intracellularly and form clusters of 32–64 spores. Both the host clone (Fi-Oer3-3) and pathogen isolate (strain 3) used in this study were sampled from Tvärminne, Finland and kept in the lab for over five years.

### Experiments overview

We conducted two microcosm experiments under laboratory conditions to determine how food provision influences host–pathogen interactions in female *D. magna*. Both experiments followed identical protocols and had the same food treatments but differed in their measurements and primary focus. In Experiment 1, *O. colligata* was grown within its host under a gradient of six different food conditions. Pathogen burden and infection status were measured after 28 days, with 16 replicates per treatment, providing a snapshot of pathogen growth. Experiment 2 consisted of two components and used the same six levels of food. The first component repeated the measurement of parasite fitness at a single time point, as in Experiment 1, but at 35 days instead of 28 (15 replicates). For the second component, 35 replicates of both pathogen-exposed and pathogen-free animals were monitored until natural death to assess effects on host survival and fecundity (the latter scored in 10 of these replicates). In addition, to determine if the rate of encounter of the host with the *O. colligata* spores depended on the food provided (as food availability may alter the host’s feeding rate), we also measured filtration rates in 10 additional juveniles per food treatment in Experiment 2. As before, we also estimated pathogen fitness by measuring infection rates and burden after animals naturally died.

## Experimental Procedure

### Host & Pathogen Preparation

To minimise maternal effects and to ensure an adequate supply of juvenile female *D. magna*, ∼100 and ∼400 adult females, for respectively Experiment 1 and 2, were held under standardised conditions for 28 days prior to the experiment. Specifically, 10-12 individuals were maintained in 400mL microcosm containing 300mL Artificial Daphnia Medium (ADaM), modified to include 5% of the recommended selenium dioxide (SeO2) concentration (Klüttgen et al., 1994). Animals were maintained under optimal growth conditions (20°C, *ad libitum supply* of batch-cultured *Scenedesmus* sp. algae grown in WC medium (Andersen et al., 2005), and were transferred to clean microcosms every four days. Prior to the start of both experiments, gravid females were transferred to new microcosms. Juveniles born within the subsequent 48 hours were collected and sexed, and only females were retained for the experiment (Figure 1). These juvenile female *Daphnia* were individually transferred into 100mL microcosms with 70mL ADaM and were randomly assigned to each of the experiment’s food treatments (3, 7, 11, 15, 19, or 23 million cells of *Scenedesmus* sp./mL every two days). A small amount of Cetyl alcohol was added to each microcosm to disrupt surface tension, mitigating potential mortality due to increased buoyancy due to small air bubbles introduced after sorting the animals by sex under the microscope. During both experiments, microcosms were kept under standard laboratory conditions (16:8 light:dark, 20**°**C) and animals were fed their respective food level every two days until the experiment’s end. Each individual was transferred to a clean microcosm every four days to avoid build-up of food and remove any offspring produced.

**Figure 1:**
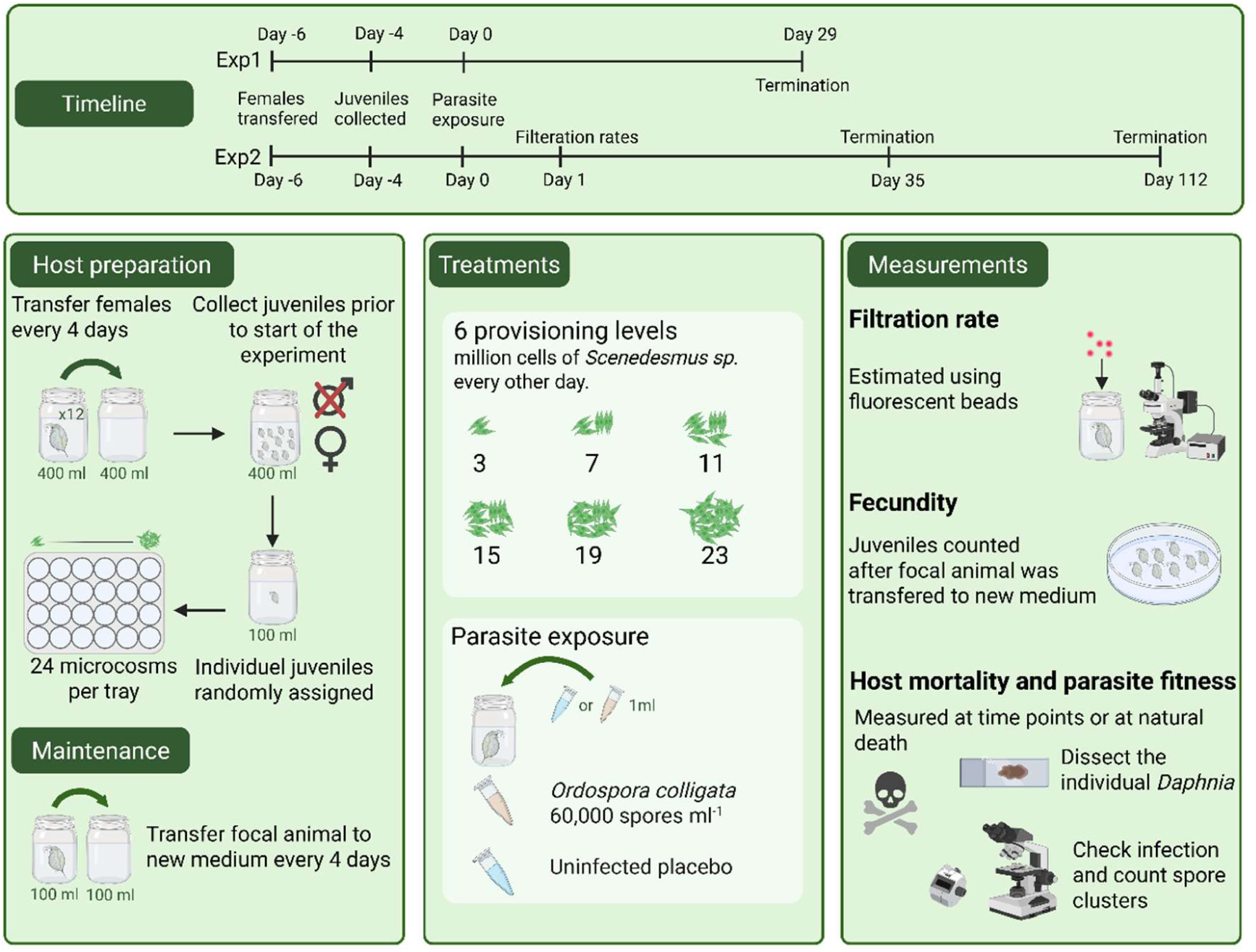
Overview of the two experiments. Both experiments were identical in terms of preparation, maintenance, and treatments, but were terminated at different times. Experiment 1 ended after 29 days, whereas in Experiment 2, animals were either dissected after 35 days or 112 days. In both experiments, host fecundity, mortality, and parasite infection rates and burden were measured. Additionally, in Experiment 1, filtration rates were measured early in the experiment.

### Pathogen Preparation & Exposure

Four days after the start of each experiment animals were either exposed to *O. colligate* or a placebo. The infectious doses were prepared by grinding infected animals (from laboratory stock cultures) with a known average pathogen burden (quantified using phase-contrast microscopy following dissection) into a slurry with a mortar and pestle. This suspension was diluted with ADaM to a concentration of ∼10,000 spores mL⁻¹, and 1 mL was added to all exposed animals across all six food treatments in both experiments. Uninfected animals in both experiments received a 1 mL placebo dose consisting of homogenized uninfected animals (Figure 1). Experiment 2 followed a fully factorial design, whereas in Experiment 1, only 12 placebo inoculations were included as a negative control.

### Measurements

*Daphnia* mortality was monitored daily. Upon natural death or termination at pre-set timepoints (Day 29 and 35), *Daphnia* were assessed for infection. The animal’s gut was dissected and inspected for the presence of *O. colligata* spore clusters under 400x magnification using bright field and phase contrast microscopy. If present, clusters of *O. colligata* were counted in the entire gut. In addition to mortality, we also recorded host fecundity by determining lifetime reproductive success. Offspring were counted by decanting any offspring that remained in the microcosm after the focal animal was transferred to a new microcosm into a large petri dish and placing this on a light box (Figure 1). Filtration rates were measured one day after exposure to *O. colligata* on a subset of 10 *Daphnia* per treatment combination (6 food levels × 2 pathogen exposure levels) to assess whether food availability directly affected encounter rates with the pathogen. Animals were transferred to a new microcosm with 35 ml of ADAM, their assigned food level and 0.52 million fluorescent microbeads (Fluoresbrite Polychromatic Red Microspheres). Animals were allowed to filter for two minutes before they were immobilized using carbonated mineral water. The number of beads in the gut was subsequently counted using a fluorescent microscope.

## Statistics

To evaluate the effects of food provisioning on pathogen and host fitness in both experiments, we used Generalized Linear Models (GLMs) implemented in R (version 4.3.1, “Beagle Scouts” (R Core Team, 2023)). One male that had been misidentified as female was omitted from the analyses for Experiment 2. For pathogen measures at days 29, 35 and 112 only those animals that died naturally or were terminated on those specific days were included in the spore burden analyses. For infectivity, all individuals that naturally died or were terminated after day 15 were included (as infection status cannot be reliably determined in early infections), except for those in which dissections failed or infection status was unclear. GLMs with a binomial error distribution were used to analyse infection rates in both experiments, with food provisioning treated as a continuous variable. GLMs were also used to analyse spore burden, host fecundity, mortality, and filtration rates; for these response variables, a negative binomial error distribution was applied as a Poisson model was over-dispersed. For all models, the best-fitting model (linear vs. quadratic vs. cubic effects of food provisioning) was selected using AIC.

## Results

### Pathogen fitness

Average infection rates were high in both experiments (70% at 29 days in Experiment 1, and 86% at 35 days and 88% at 112 days in Experiment 2) and were positively correlated with the amount of food provided. Higher food provisioning to the host for 29 and 35 days after exposure to *O. colligata* led to higher rates of infection (Table 1A, Figure 2A, 2B and P = 0.0001, P = 0.0224, respectively). For example, infection rates rose from 27% with 3 million algae to 94% when 23 million algae were provided. No relationship was found after 112 days in Experiment 2 (Table1A, Figure 2C, P = 0.2315). However, combining infection data from both time points in Experiment 2 did show an overall positive correlation with the food supply (combined 35-day and 112-day data, P= 0.01198). Similarly, the burden of *O. colligata* increased with greater provisioning. Animals dissected after 29 and 35 days showed greater burden with higher levels of food provisioning (Table 1B, Figure 2D,2E, P = 0.0003 and P<0.0001, respectively). Specifically, the number of spore clusters increased from 4 to 384 and from 110 to 493 with increasing food provision, after 29 and 35 days, respectively. A similar trend (P= 0.0759) was observed in the pathogen burden after 112 days, when the number of spore clusters rose from 655 at 3 million algae to 1880 at 23 million algae (Table 1B, Figure 2F). Moreover, when all natural deaths were included, rather than only animals sacrificed after 112 days, burden increased from 112 to 466 spores from the lowest to the highest provisioning level (P > 0.001).

**Figure 2.**
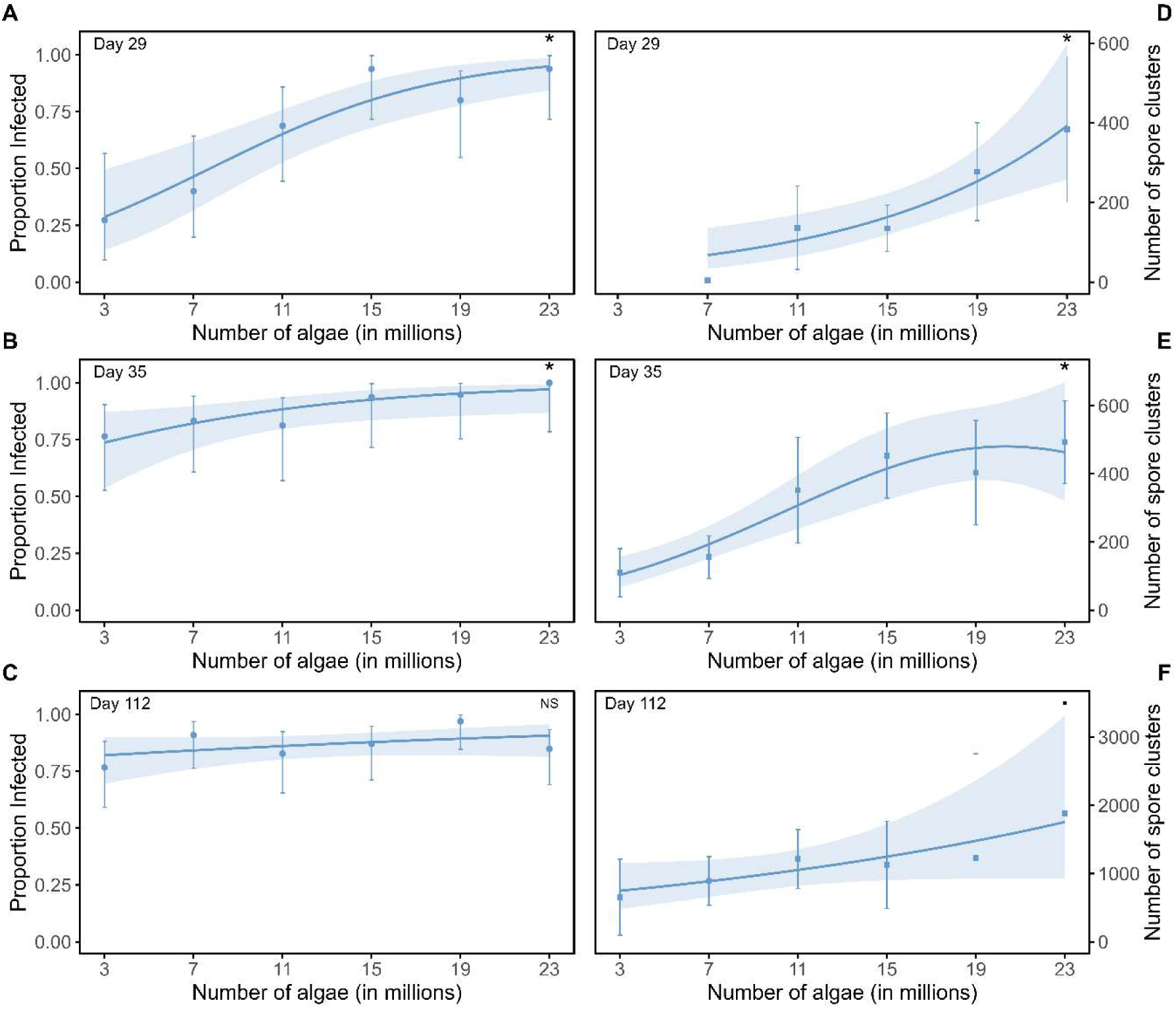
Increased food supply to the host resulted in higher pathogen fitness. Infection rates (left panels) were recorded at 29, 35, and 112 days post-exposure (Panels A, B, and C, respectively). pathogen burden, measured as the number of O. colligata spore clusters in the host gut (right panels), was also recorded at 29, 35, and 112 days post-exposure (Panels D, E, and F, respectively). Points represent the means of the food treatments; the line is the model fit and error bars and shaded areas indicate 95% confidence intervals. Symbols in the top right corner indicate significance; full stats can be found in Table 1.

**Table 1:** Generalized linear models for the effect of food provisioning on infection (binomial distribution) and burden (negative binomial distribution). Shown are estimates, standard errors (Std. Error), z-values, and p-values for the endpoints of Experiment 1 (29 days) and Experiment 2 (35 and 112 days). Linear fits had the lowest AIC for all models except for burden on day 35 were a quadratic function fit best.

| **A: Infection** |  | Estimate | Std. Error | z value | p value |  |
| --- | --- | --- | --- | --- | --- | --- |
| Day 29 (exp1) | Intercept | -1.486 | 0.578 | -2.571 | 0.0102 | * |
|  | Food | 0.192 | 0.047 | 4.045 | 0.0001 | *** |
| Day 35 (exp2) | Intercept | 0.657 | 0.579 | 1.134 | 0.2569 |  |
|  | Food | 0.125 | 0.055 | 2.283 | 0.0224 | * |
| Day 112 (exp2) | Intercept | 1.405 | 0.434 | 3.235 | 0.0012 | ** |
|  | Food | 0.038 | 0.032 | 1.196 | 0.2315 |  |
| **B: Burden** |  | Estimate | Std. Error | z value | p value |  |
| Day 29 (exp1) | Intercept | 3.452 | 0.556 | 6.21 | 0.0000 | *** |
|  | Food | 0.110 | 0.031 | 3.589 | 0.0003 | *** |
| Day 35 (exp2) | Intercept | 5.725 | 0.087 | 65.677 | <0.0001 | *** |
|  | Food (linear) | 4.185 | 0.751 | 5.574 | <0.0001 | *** |
|  | Food (quadratic) | -1.717 | 0.750 | -2.289 | 0.0221 | * |
| Day 112 (exp2) | Intercept | 6.490 | 0.285 | 22.795 | <0.0001 | *** |
|  | Food | 0.043 | 0.024 | 1.775 | 0.0759 | . |

### Host fitness

Exposure to *O. colligata* and varying levels of food availability altered host fecundity and filtration rates, while host survival remained unaffected. Greater food provisioning led to increased fecundity (Table 2, Figure 3A, P>0.0001), whereas *Daphnia* exposed to the pathogen exhibited lower fecundity compared to controls (P<0.0001). Reductions in fecundity due to the pathogen exposure ranged from 10% to 40%, with the greatest reductions observed at intermediate food levels (non-overlapping 95% confidence intervals), and smaller, non-significant reductions at both low and high food levels (overlapping 95% confidence intervals). A trend in the interaction term (Exposure × Food [quadratic], P = 0.0560) also suggests that fecundity in control and exposed individuals may respond differently to food provisioning. While pathogen exposure and food level influenced offspring production, survival remained unchanged (Table 2, Figure 3B, P = 0.129 and P= 0.782 for pathogen exposure and food provisioning, respectively), with an average lifespan of 86.5 days across all treatments. Low food levels combined with pathogen exposure led to increased filtration rates (Figure 3C, interaction of food and the combined polynomial, P =0.0203). The number of beads found in the guts of exposed animals that were fed 3 million algae was fourfold higher than in control animals, increasing from an average of 675 to 2657 beads filtered per hour.

**Figure 3:**
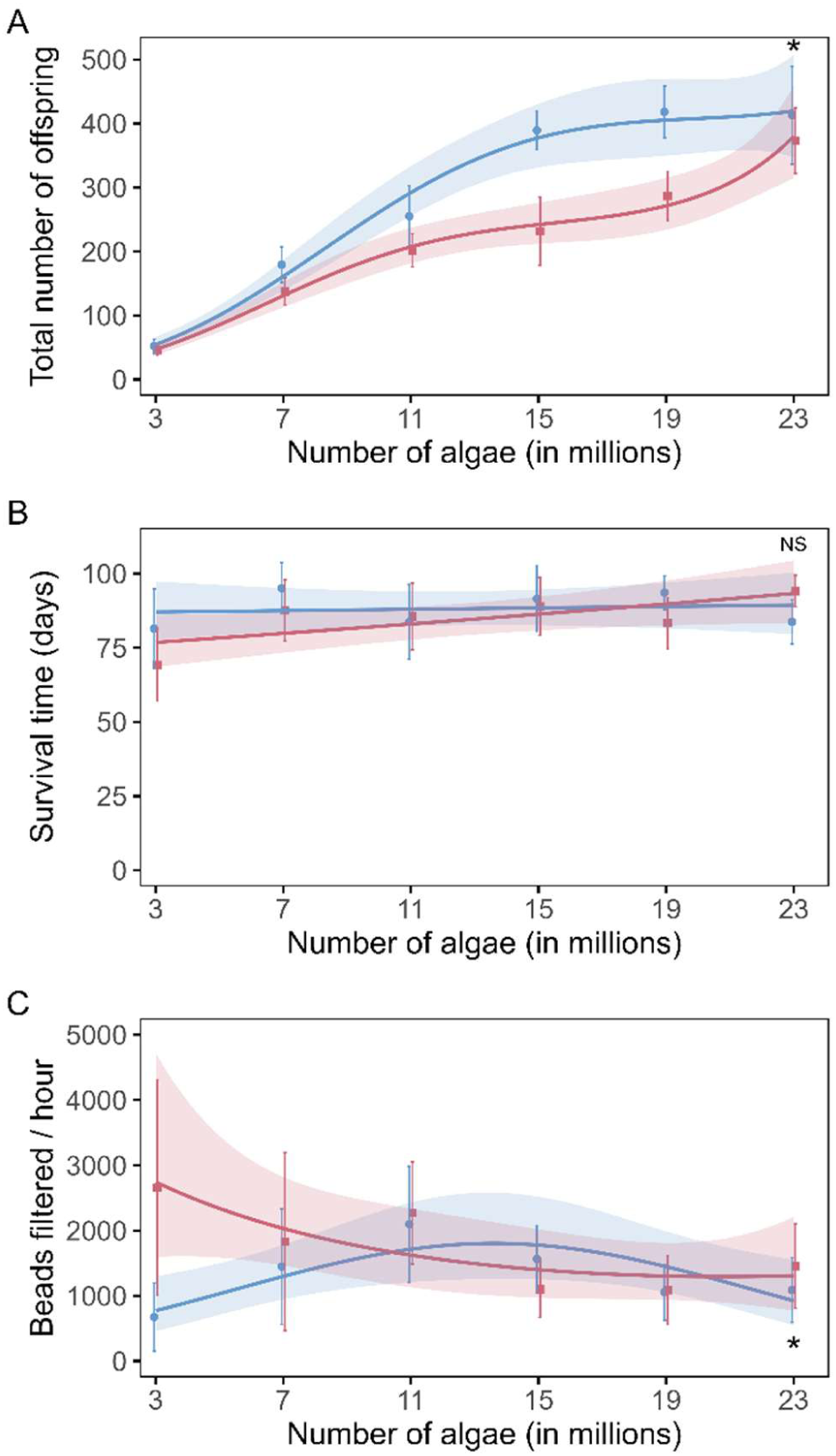
Increased food supply and exposure to O. colligata influenced hosts’ fecundity and filtration but not survival. Total lifetime reproductive success at the end of the experiment (panel A), survival at the end of the experiment (panel B) and filtration rates of juveniles (panel C). Points represent the means of the food treatments; the line is the model fit, and error bars and shaded areas indicate 95% confidence intervals. The animals exposed to O. colligata are in red, and the unexposed control Daphnia in blue. Symbols in the top right corner indicate significance; full stats can be found in Table 2.

**Table 2:** Generalized linear models using a negative binomial distribution for the effect of food provisioning and exposure to O. colligata on D. magma fecundity, survival and filtration rate. Shown are estimates, standard errors (Std. Error), z-values, and p-values for the endpoint of Experiment 2 (112 days). A cubic, linear, and quadratic fit provided the best model fit (lowest AIC) for fecundity, survival, and filtration rate, respectively.

| Host traits (Experiment 2, day 112) |  |  |  |  |  |  |
| --- | --- | --- | --- | --- | --- | --- |
| Fecundity | Intercept | 5.456 | 0.041 | 133.197 | <0.0001 | *** |
|  | Food (linear) | 7.074 | 0.455 | 15.560 | <0.0001 | *** |
|  | Food (quadratic) | -3.583 | 0.452 | -7.926 | <0.0001 | *** |
|  | Food (cubic) | 0.906 | 0.448 | 2.020 | 0.0433 | * |
|  | Exposure | -0.274 | 0.058 | -4.715 | <0.0001 | *** |
|  | Exposure * Food (linear) | -0.232 | 0.645 | -0.360 | 0.7190 |  |
|  | Exposure * Food (quadratic) | 1.226 | 0.642 | 1.911 | 0.0560 | . |
|  | Exposure * Food (cubic) | 0.673 | 0.636 | 1.058 | 0.2902 |  |
| Survival | Intercept | 4.463 | 0.069 | 64.426 | <0.0001 | *** |
|  | Food | 0.001 | 0.005 | 0.277 | 0.782 |  |
|  | Exposure | -0.149 | 0.098 | -1.518 | 0.129 |  |
|  | Food* Exposure | 0.008 | 0.007 | 1.238 | 0.216 |  |
| Filtration rate | intercept | 7.140 | 0.119 | 59.838 | <0.0001 | *** |
|  | Food (linear) | 0.609 | 1.290 | 0.472 | 0.6369 |  |
|  | Food (quadratic) | -3.24 | 1.28 | -2.526 | 0.0115 | * |
|  | Exposure | 0.2799 | 0.172 | 1.63 | 0.1031 |  |
|  | Exposure * Food (linear) | -3.324 | 1.849 | -1.797 | 0.0723 | . |
|  | Exposure * Food (quadratic) | 4.268 | 1.850 | 2.307 | 0.021 | * |

## Discussion

Food provisioning shaped host filtration rates, pathogen performance, and host fitness, but effects were non-linear and trait-specific. Pathogen fitness increased with greater food availability, consistent with the idea that pathogens can be resource-limited (Clasen and Elser, 2007). This pattern is also consistent with theory predicting that increased resource supply benefits pathogens when resource use by the pathogen and host immune system is independent (Cressler et al., 2014). In contrast, host fitness responses did not scale uniformly with food supply: survival remained unchanged, whereas infection-related fitness costs (virulence) peaked at intermediate provisioning levels. At low food provisioning, reduced pathogen burden limited host damage, while at high food levels, hosts may have better tolerated infection despite higher pathogen loads. Together, these results show that food availability simultaneously modifies multiple host and pathogen traits, producing complex outcomes for virulence.

Filtration rates were highest in exposed animals with limited access to food. A negative correlation between food provisioning and feeding rates has been reported previously for *Daphnia* (Chow-Fraser and Sprules, 1992) and other organisms (Liu et al., 2010). However, why exposed and control animals differ in their filtration rates is less clear. Damage to gut cells caused by invading *O. colligata* spores may reduce the efficiency of the digestive tract and raise feeding rates. Alternatively, pathogen-exposed animals may have higher nutrient demands due to the cost of immune defences (Lochmiller and Deerenberg, 2000; Schmid-Hempel, 2005) or as a consequence of a general stress response (Sokolova, 2013). Moreover, because spores of *O. colligata* are encountered during feeding (Ebert, 2005), higher filtration rates at low food levels would be expected to lead to greater disease prevalence. However, the opposite was observed; the lowest level of food provisioning had the lowest infection rates.

Pathogen fitness increased with higher food provisioning. Because of their rapid growth, pathogens often have different nutrient requirements than their hosts (Elser et al., 2003) and may be limited when food availability or nutrient quality is low. For example, growth of a virus of the freshwater alga *Chlorella* was strongly reduced when phosphorus availability was low (Clasen and Elser, 2007). Moreover, theory shows that increased pathogen fitness with higher levels of food either occurs if there is no competition for resources between the immune system and the pathogen or when the pathogen can restrict the action of the immune system (Cressler et al., 2014). Although *Daphnia* shows variation in immune responses such as phenol oxidase and haemocyte activity (Auld et al., 2010; Mucklow et al., 2004), how this variation translates into resistance (Ebert et al., 2016), and whether pathogens compete with the host immune system for resources, remains unclear for many of its pathogens. However, because immunity in *Daphnia* can be energetically costly (that is costs of resistance have been reported (Dallas et al., 2016; Hall et al., 2024)), *O. colligata*, as a gut pathogen, may gain early access to host resources, potentially restricting the energy available for immune defence. Increased food quantity or quality has been shown to elevate pathogen burden in Daphnia, including for the gut microsporidian *Glugoides intestinalis* (Pulkkinen and Ebert, 2004)) and the fungal pathogen *Metchnikovia bisporate* (Fearon et al., 2025; Hall et al., 2009). Similarly, the microsporidia *Binucleata daphniae* and *O. colligata* exhibit increased fitness under nutrient-enriched conditions (Decaestecker et al., 2015). Indeed, enhanced food quality in one generation may even alter disease burden in subsequent populations through maternal effects (Schlotz et al., 2013). However, effects are not universal as no correlation between food and pathogen performance were found in the *Daphnia* bacterical pathogen *Pasteuria ramosa* (Fearon et al., 2025; Schoebel et al., 2014). In general, many invertebrate systems have reported an increase in pathogen fitness under food supplementation (Bedhomme et al., 2004; Cooper et al., 2009; Hall et al., 2009) while vertebrate systems have often reported the opposite (Edirisinghe et al., 1981; Shlomai et al., 2006). Pathogens of small-bodied invertebrate hosts could be limited by resource availability and greater food supply may lead to increasing growth, as opposed to vertebrates, which might be immune-limited (Cressler et al., 2014). While *O. colligata* fitness increases with greater food provisioning, food availability can generate non-linear effects by affecting multiple life history traits at once, which can lead to complex host–pathogen interactions (Borer, Kendig & Holt, 2023).

Host fitness scaled non-linearly with food provisioning, and different fitness traits were affected in distinct ways. Survival was not influenced by either food provisioning or pathogen exposure. However, infection-related fitness costs (i.e., virulence) were greatest at intermediate levels of food provisioning. Food provisioning simultaneously alters numerous host and pathogen fitness traits (e.g., host fecundity, mortality and feeding behaviour and pathogen growth and transmission)(Borer et al., 2023), and because effects on R₀ (the basic reproductive number of a disease), disease dynamics, and virulence depend on the interplay among these traits, changes in food quality and quantity can lead to complex interactions. For example, in plants infected with the necrotrophic fungal pathogen *Botrytis cinerea*, nitrogen supply is linked to both plant defences and the expression of virulence genes in the pathogen, resulting in disease severity being highest at both low and high levels of nutrient availability (Abro et al., 2013). In *Daphnia*, low food levels reduced pathogen burden, resulting in minimal impact on the host. At higher food levels, pathogen abundance increased, leading to greater damage to the host and reduced fecundity. At the highest provisioning levels, however, hosts may have had sufficient energy reserves to compensate for the increased pathogen load and were therefore better able to tolerate infection, leading to higher fecundity. Increased tolerance despite unchanged or even increased pathogen burdens with greater resource availability has also been found in the burying beetle, *Nicrophorus vespilloides* infected with *Photorhabdus luminescens* (Miller and Cotter, 2018) and great tits, *Parus major* exposed to the flea *Ceratophyllus gallinae* (see for a review Budischak and Cressler, 2018; Christe et al., 1996). Moreover, that virulence can be highest at intermediate levels of provisioning has also been seen before *Aedes aegypi* musquitos infected with *Vavraia culi* (Bedhomme et al., 2004). However, scaling of virulence with provisioning is diverse, in part due to the complex interplay of multiple traits (Borer et al., 2023), and increased and decreased virulence with food provisioning has also been frequently reported (see Pike et al., 2019 for a review).

Virulence is an emergent property arising from interactions between hosts and their pathogens and can, among others, be influenced by host resource acquisition, pathogen growth, and host tolerance. Indeed, in our system, a large range of virulence has been reported (from negligible impact to 60% reduction in fecundity (Kirk et al., 2018; McCartan et al., 2025; O’Keeffe et al., 2024)). While *O. colligata* is often described as a benign pathogen (Ebert, 2005), we show that variation in resource availability can alter host tolerance and pathogen growth in ways that may increase virulence. Our findings add to a growing body of work demonstrating that resource availability shapes not only resistance but also tolerance (Christe et al., 1996; Miller and Cotter, 2018). However, significant gaps remain: empirical studies report opposing effects of resources on tolerance, and theory explaining these discrepancies requires further development, particularly in the context of ongoing environmental change (see, Budischak and Cressler, 2018 for a review on resource availability and tolerance). Indeed, with eutrophication and anthropogenic resource enrichment, resource shifts may reshape host–pathogen interactions and alter disease dynamics (Cárdenas et al., 2018; Johnson et al., 2010), which can have far-reaching impacts. For example, in *Daphnia*, a disease sensitive to food provisioning (Fearon et al., 2025; Hall et al., 2009) has been shown to induce trophic cascades (Duffy, 2007).

Moreover, with food availability affecting pathogen virulence, changing levels of provisioning may either amplify or decrease the strength of selection pathogens and parasites may place on their host which can impact long term evolution of resistance and tolerance (Zeller and Koella, 2017), alter the outcome of multiple infection (Budischak et al., 2015) and impact disease dynamics though eco evolutionary feedback loops (Hall et al., 2009). Indeed, by demonstrating that resource availability can convert a seemingly benign pathogen into a more virulent one, our study underscores how ongoing environmental change may alter disease outcomes.

## Acknowledgements

The authors thank all laboratory members for their support with the empirical work for their direct assistance during the experiment. The authors thank Dieter Ebert and Jürgen Hottinger for provision of the *D. magna* genotypes.

## Author’s contribution

PL, NZ and JP developed the initial research question and PL, NZ, JP CCB and MO contributed to the design of the experiment. NZ and PL conducted the first bloc of the experiment. CCB and MO led the second experimental bloc, with the assistance of NMcC, JS and PL. CCB and PL conducted the analyses and generated figures. CCB, MO and PL wrote the first draft of the manuscript and all authors contributed to revisions. All authors read and approved the final manuscript.

